# Sequential restriction of SNARE-mediated fusion by the COPII inner coat and SNARE chaperones

**DOI:** 10.64898/2026.02.02.703378

**Authors:** E.J. Mackey, T. Takenaka, E. A. Miller, A.J. Merz

## Abstract

SNARE-mediated fusion requires assembly of four SNARE domains (R, Qa, Qb, and Qc) distributed across two membranes, with an SM (Sec1/Munc-18 family) chaperone catalyzing this assembly. We investigated topological requirements for the four SNAREs that mediate ER-Golgi fusion using *in vitro* assays. In the presence of the cognate SM Sly1, we find that a single topology drives efficient fusion: the R SNARE on one membrane versus Qa/Qb/Qc SNAREs on the other. These results prompted us to look upstream, at COPII-SNARE interactions. The assembled COPII coat is known to block fusion. We discovered that of five COPII core subunits, the Sar1 GTPase and Sec23/Sec24 subunits were necessary and sufficient to prevent fusion for prolonged periods, and that Sar1 was dispensable for inhibition over short periods. When Sec24- interactions with vesicle SNAREs were disrupted, inhibition was relieved, indicating that SNARE sequestration by COPII prevents fusion. These observations help to explain how appropriate fusion events are facilitated while inappropriate events are deterred.

## Introduction

Vesicular traffic within the endomembrane system controls subcellular protein distribution, organelle biogenesis and turnover, and secretion. The final step of vesicular transport, fusion of two compartments, is a nexus of specificity and regulation. Fusion events are executed by the SNARE (Soluble N-ethylmaleimide-sensitive factor attachment protein receptor) protein family (Bombardier and Munson, 2015; Jahn et al., 2023). Fusion requires assembly of four distinct SNAREs into a parallel α-helical bundle that spans the donor and acceptor membranes — the pre-fusion *trans* SNARE complex (Hanson et al., 1995; Poirier et al., 1998; Sutton et al., 1998; Weber et al., 1998). Following fusion, the hyperstable *cis* SNARE complex is disassembled by an ATPase complex composed of Sec17 and Sec18 (α-SNAP and NSF, in mammals). This recycles and reenergizes SNAREs so that they can catalyze subsequent rounds of fusion (Hanson et al., 1995; Khan et al., 2025; Mayer et al., 1996).

Each class of membrane fusion event employs a unique set of SNAREs, though some SNARE proteins operate in more than one pathway (Grissom et al., 2020). Each SNARE bundle contains four parallel helices in a stereotypical arrangement (Fasshauer et al., 1998; Sutton et al., 1998). Based on sequence homology and position within the bundle, the four helices are named Qa, Qb, Qc, and R. In this nomenclature, Q and R denote the presence of either glutamine (Q) or arginine (R) residues that contribute side chains to a central “ionic” packing layer near the middle of the bundle. At the presynaptic membranes of neurons, only two SNAREs are anchored by C-terminal transmembrane helices: VAMP2/Synaptobrevin (R) resides on vesicles, whereas Syntaxin1a (Qa) is on the plasma membrane. A third SNARE, SNAP-25, contains both Qb and Qc helices, separated by an unstructured linker. SNAP-25 associates with the target membrane through palmitoyl modifications to the linker (Rizo and Xu, 2015). Most fusion events, however, employ four separate SNARE polypeptides, each bearing its own C-terminal membrane anchor. Defining which SNAREs must reside on donor versus acceptor membranes is the SNARE topology problem.

SNAREs bearing C-terminal transmembrane domains are inserted into membranes at the ER through the tail-anchor pathway (Mateja and Keenan, 2018). SNAREs are then exported from the ER as cargo in COPII vesicles (Miller et al., 2002). They then traffic to steady-state locations in the secretory and endolysosomal systems. These locations are dynamically maintained through cycles of anterograde and retrograde traffic. During these processes, there are numerous opportunities for off-target fusion to occur. Consequently, mechanisms that restrict the range of allowed SNARE fusion topologies are thought to be critical determinants of directionality and specificity — and ultimately, of organelle identity (Koike and Jahn, 2022). The sets of SNARE topologies that can assemble into productive fusion machines *in vivo* are constrained by: (*i)* the intrinsic assembly properties of SNAREs themselves (McNew et al., 2000a); (*ii)* chaperones that catalyze *trans*-SNARE complex assembly and proofread assembled complexes (Choi et al., 2018; Dubuke and Munson, 2016; Schwartz et al., 2017; Zhang and Hughson, 2021); and (*iii*) factors such as vesicle coats that control which SNARE proteins are present on which membranes (Miller et al., 2002; Mossessova et al., 2003).

Among the SNARE chaperones, <u>S</u>ec1/<u>m</u>ammalian Unc-18 (SM) proteins are central. SMs are thought to be essential for all SNARE-mediated fusion in cells (Rizo and Sudhof, 2012). After many years of controversy, breakthrough structures of the endolysosomal SM Vps33 suggested a unifying model: SMs are enzymes that catalyze assembly of a half-zipped *trans* “template complex” between the Qa and R SNAREs (Baker et al., 2015; Jahn et al., 2023; Rodkey et al., 2008). This model makes a key prediction about SNARE topology: Qa and R SNAREs must always function *in trans* to one another. How SM proteins interact with Qb and Qc SNAREs, and whether SMs impose additional topological restrictions on Qb or Qc SNARE function, are unresolved questions.

We study the first trafficking event in the secretory pathway: fusion of ER-derived COPII carriers at the *cis-*Golgi or the precursor ER-Golgi intermediate compartment (ERGIC). The COPII coat comprises five subunits that sequentially assemble at the ER exit site (Antonny et al., 2001; Sato and Nakano, 2005). Upon GTP binding, the Sar1 GTPase associates with the membrane (Van der Verren and Zanetti, 2023). Sar1 then recruits the Sec23/24 complex (or variant complexes containing Sec24 paralogs). Sec24 binds the cytoplasmic tails of transmembrane cargo molecules to capture and concentrate them for ER export. Finally, the Sec13/31 complex associates with and stabilizes the nascent coat lattice. Although the spatiotemporal dynamics of COPII uncoating and the features of COPII carriers are under intense discussion, uncoating cannot occur until Sar1-bound GTP is hydrolyzed. Before GTP hydrolysis, COPII blocks fusion of the carrier with target membranes (Barlowe et al., 1994; Cai et al., 2007b; Oka and Nakano, 1994). Though the mechanism through which COPII blocks fusion has not been fully dissected, COPII contains SNARE binding sites that might prevent SNAREs from entering pre-fusion complexes (Mossessova et al., 2003; Miller et al., 2003). *In vivo*, COPII may uncoat partially, or in stages. COPII subcomplexes may contribute to vesicle tethering at Golgi precursor membranes (Angers and Merz, 2011; Cai et al., 2007a).

There are seven possible 1:3 or 2:2 SNARE topologies. In yeast, COPII-mediated anterograde traffic requires four SNAREs: Sed5 (Qa), Bos1 (Qb), Bet1 (Qc), and Sec22 (R) (Banfield et al., 1994; Newman and Ferro-Novick, 1987; Novick et al., 1980; Shim et al., 1991; Sogaard et al., 1994). At least three different — and incompatible — topologies have been proposed to catalyze fusion in the COPII pathway (Fig. 1E, 1C or 1F, and 1I). In studies with chemically defined reconstituted proteoliposomes (RPLs), Rothman’s lab assigned Bet1 (Qc) to the COPII vesicle, opposite Sed5 (Qa), Bos1 (Qb), and Sec22 (R) on the target membrane (McNew et al., 2000a; Parlati et al., 2000; Parlati et al., 2002). However, no SM chaperone was present in these fusion experiments. Work using complex yeast cell extracts and temperature-sensitive SNARE mutants challenged the Rothman lab’s interpretation (Cao and Barlowe, 2000; Liu and Barlowe, 2002; Miller et al., 2005). These experiments suggested that Sec22 (R), Bos1 (Qb) and Bet1 (Qc) function on the COPII vesicle, opposite Sed5 (Qa) on the Golgi target membrane. Experiments using chemically defined RPLs, but now in the presence of the SM Sly1, yielded yet another result: robust fusion was observed with the R SNARE opposite Qa, Qb, and Qc (Duan et al., 2024a; Duan et al., 2024b; Furukawa and Mima, 2014; Jun and Wickner, 2019). These studies, however, did not explore alternative SNARE topologies.

**Figure 1.**
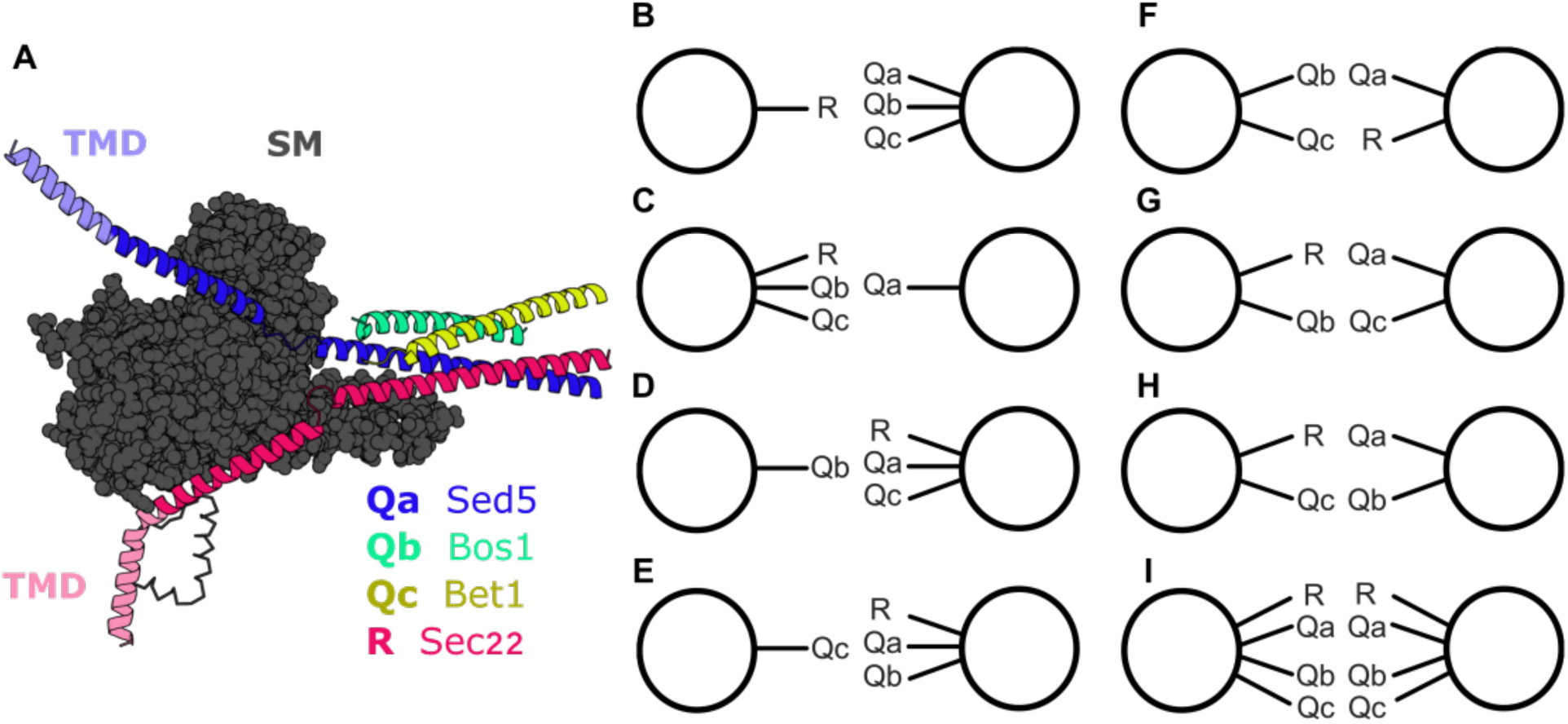
Possible SNARE topologies. (**A**) AlphaFold 3 model of the ER-Golgi trans-SNARE bundle in a template complex with Sly1. Qa and R SNAREs are modeled including only their SNARE domains through their transmembrane helices (Qa: 235-340, R: 111-214). The transmembrane domains are depicted in a lighter shade, as noted. Qb (residues 149-182) and Qc (residues 36-78) SNARE domains are modeled from α-helical packing layers −8 to −3 within the SNARE bundle (Fasshauer et al., 1998; Sutton et al., 1998). (**B-I**) Schematics represent the RPL pairs prepared and tested. (**B-H**) The seven minimal, unique SNARE topology pairs are shown as well as, (**I**) the eighth topology tested, which mimics homotypic fusion.

Here, we use chemically defined reconstitutions to explore all possible ER-Golgi SNARE topologies, in the presence and absence of the essential SNARE chaperones Sly1 (SM), Sec17, and Sec18. In the absence of Sly1 there is little or no fusion, consistent with the lethal phenotype of *sly1Δ* null alleles. In the presence of Sly1, one of seven possible topologies catalyzes efficient fusion: R opposite Qa, Qb, and Qc. Building on these results, we explore how the COPII coat prevents fusion until the coat depolymerizes. Surprisingly, although COPII packages all four SNARE proteins into ER-derived carriers, we find that COPII-SNARE binding, through a specific site on Sec24, is necessary to block fusion. We propose a working model in which both COPII suppression of fusion, and Sly1 facilitation of fusion, are consistent with a unique functional SNARE topology. This arrangement may not only facilitate anterograde traffic, but also provide a mechanism for suppressing inappropriate fusion events across alternative topologies that might disrupt organelle identity and maturation.

## Results

### Sly1 stimulates fusion with strict SNARE topological restrictions

Crystal structures of SNAREs complexed with the lysosomal SM Vps33 suggest that R and Qa SNAREs must function *in trans* (Baker et al., 2015). This is consistent with the arrangement of neuronal exocytic SNAREs, which have only two rather than four transmembrane anchors. Elegant force spectroscopy studies corroborated this model, demonstrating that an SM, a Qa-SNARE, and an R-SNARE can form a partially-zipped intermediate called the template complex (Baker et al., 2015) of SM proteins (Sly1, Sec1, and Vps45) with client SNARE complexes implies that the requirement for Qa and R-SNAREs to act on opposite membranes is generalizable (Fig. 1A; Fig. S1; Fig. S2)(Baker et al., 2015). We therefore hypothesized that *any* arrangement that places the R and Qa SNAREs *in trans* should drive fusion in the presence of a cognate SM. Accordingly, Qb and Qc SNAREs should be able to drive fusion when anchored in either membrane.

To test this model, we prepared RPLs bearing all possible topologies: four sets with one SNARE versus the remaining three (Fig. 1B-E), three sets with pairs of two SNAREs in opposition (Fig. 1F-H), and one set with all four SNAREs present on both membranes, recapitulating homotypic vesicle fusion (Fig. 1I). The SNAREs and chaperones were present at near-physiological concentrations (Supplementary Table S1). Of the seven unique topologies, four place the R and Qa SNAREs *in trans*, while three place the R and Qa *in cis*. We measured fusion between RPL populations using two orthogonal FRET pairs, one reading out lipid mixing and the other, content mixing. Our analyses focused on content mixing, the fusion reaction endpoint. We set up reactions with paired RPL populations, took baseline fluorescence readings, and then added Sly1 to initiate fusion. To our surprise, Sly1 initiated rapid and efficient fusion with only one of the seven unique topologies: the R SNARE versus the Qa/Qb/Qc SNAREs (Fig. 2A-B). With this topology rapid fusion immediately followed Sly1 addition.

**Figure 2.**
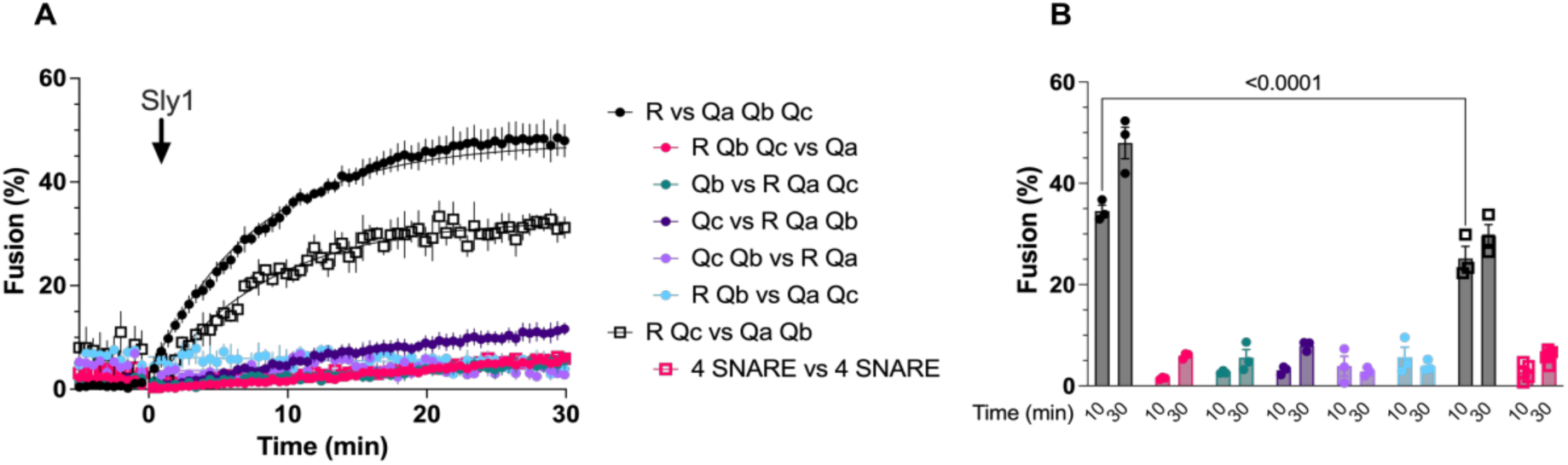
Rapid, efficient fusion in the presence of Sly1 occurs with select topologies (**A**) Fusion assays were carried out for each pair of RPLs. Fusion reactions ran for thirty minutes after initiation. RPL populations were mixed together for a baseline reading. Reactions were initiated by the addition of 100 nM Sly1. Lines show fits of a 2-parameter kinetic model. 100% fusion was determined by determinig the total content mixing of two RPL populations by bursting with detergent. (**B**) Fusion for each configuration at 10 and 30 min. is shown, from the datasets plotted in (A). Both plots show means from 3 separate experiments, with error bars spanning ± S.E.M.). The P value indicated by the horizontal bar in (B) is from 2-way ANOVA with correction for multiple comparisons.

A second topology, R/Qc vs. Qa/Qb, drove substantial fusion but with slower kinetics. Ten minutes into the reaction, about half as much fusion occurred as with the R vs. Qa/Qb/Qc topology (Fig. 2B). Thus, there appears to be some plasticity for Qc SNARE topology; however, the lower efficiency of the R/Qc opposite Qa/Qb topology suggests that it is not preferred. In extensive mutational studies of Sly1 and the Qa-SNARE Sed5, even a ∼50% decrease in fusion *in vitro* correlated with conditional lethality *in vivo* (Duan et al., 2024a; Duan et al., 2024b). Among the seven unique topologies only these two drove substantial fusion, with the remaining five virtually inactive.

These findings are in tension with previous work suggesting that fusion is catalyzed solely by the Qc vs. R/Qa/Qb topology (McNew et al., 2000a; Parlati et al., 2000; Parlati et al., 2002). That conclusion was based on lipid mixing experiments in a chemically defined system, where Sly1 was not present. The apparent discrepancy results, at least in part, from our use of content mixing as a primary endpoint. When we examined lipid rather than content mixing, very slow but reproducible lipid mixing was detected with the the Qc vs. R/Qa/Qb topology. This signal was independent of Sly1 addition. In comparison, the R vs. Qa/Qb/Qc topology catalyzed essentially no lipid mixing unless Sly1 was present, at which point the signal quickly outpaced the Qc vs R/Qa/Qb topology (Fig. S3). Lipid mixing without content mixing could indicate accumulation of a stalled or hemifused state. Thus, our experimental findings are at least partially consistent with the Rothman group’s. Nevertheless, as content mixing is minimal and unchanged in the presence of Sly1, and given that loss of Sly1 is lethal *in vivo* (Duan et al., 2024b; Ossig et al., 1991), we suggest that the Qc vs. R/Qa/Qb and the R/Qc vs Qa/Qb topologies do not catalyze physiologically relevant fusion *in vivo*.

We also tested RPLs with all four SNAREs on both membranes, a configuration that mimics homotypic fusion, as reported to occur with mammalian COPII carriers in a cell-free system (Xu and Hay, 2004). These events require the mammalian Sec17/Sec18 homologs ɑ-SNAP/NSF, ATP, and mammalian Sly1 (SCFD1). As expected from these studies and experiments with RPLs (Duan et al., 2024a; Duan et al., 2024b; Jun and Wickner, 2019), in the absence of Sec17/18:ATP we observed little or no homotypic fusion of 4-SNARE RPLs (Fig. 2A-B). Thus, we next asked whether Sec17/18:ATP can promote Sly1-dependent fusion across a broader range of SNARE topologies.

### Sec17/18 and SNARE topology

Sec17/18:ATP disassemble post-fusion *cis-*SNARE complexes, releasing SNAREs to catalyze subsequent rounds of fusion (Hanson et al., 1997; Mayer et al., 1996). These enzymes also disassemble unproductive *cis-* and *trans-*SNARE complexes that assemble aberrantly (Choi et al., 2018; White et al., 2025), and they can in some cases help otherwise non-fusogenic SNARE complexes to function. We hypothesized that cycles of SNARE assembly and disassembly might allow alternative SNARE topologies multiple chances to form productive *trans* complexes, leading to fusion. Alternatively, Sec17 and Sec18 might directly stimulate SM-dependent fusion (Ma et al., 2016; Orr and Wickner, 2022; Schwartz et al., 2017; Schwartz and Merz, 2009; Song et al., 2021; Zick et al., 2015).

In the presence of Sec17/18:ATP and Sly1, again at near-physiological concentrations, there was rapid fusion with the 4 SNARE vs 4 SNARE (homotypic) configuration (Fig. 3H), consistent with previous work. No fusion occurred when R and Qa SNAREs opposed Qb and Qc SNAREs (Fig. 3E). Some fusion did occur when the R/Qc pair opposed Qa/Qb (Fig. 3G), but this fusion was slower than with the R vs. Qa/Qb/Qc topology, and Sec17/18 and ATP did not increase the rate or total amount of fusion for this topology.

**Figure 3.**
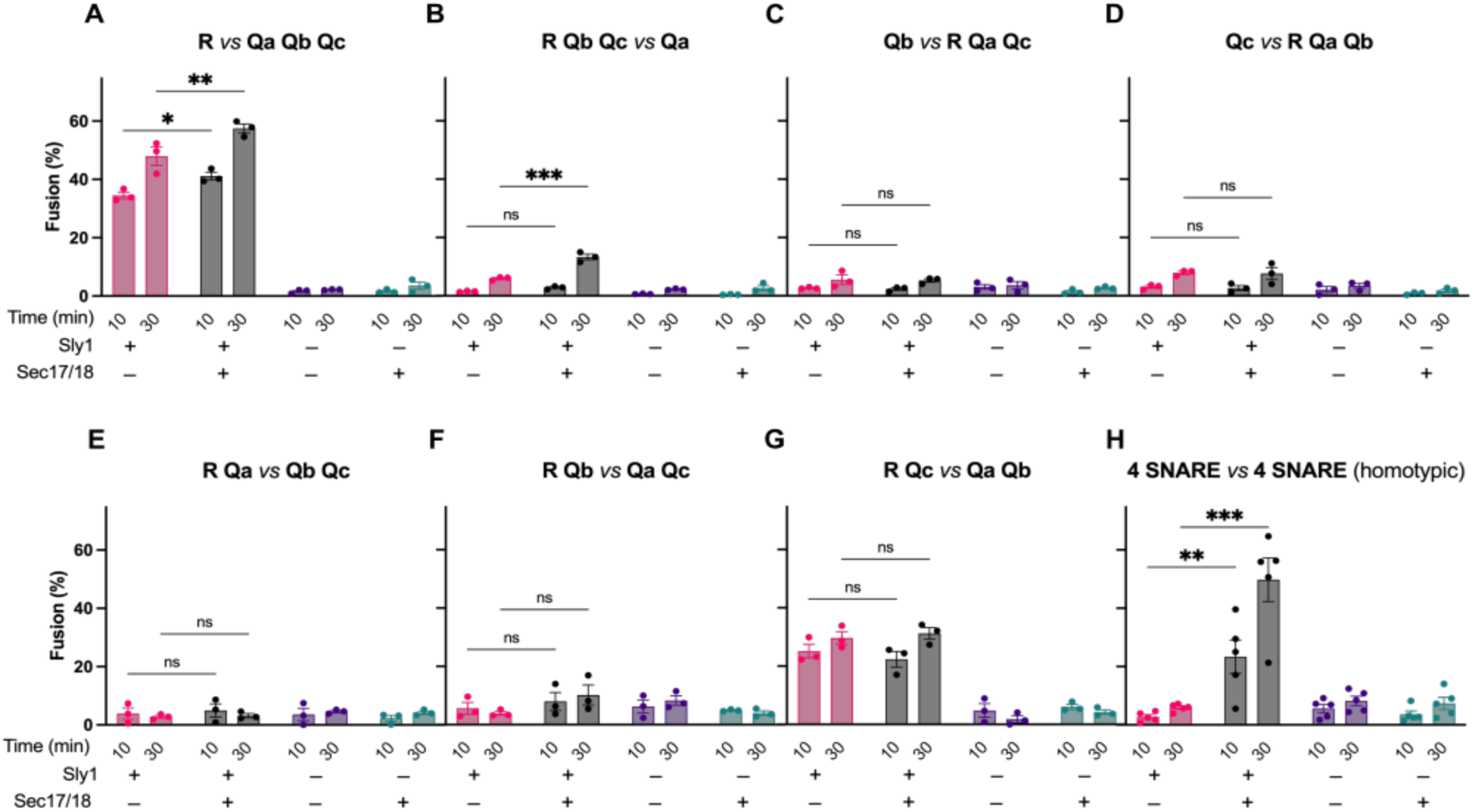
Sec17/18 and topological restriction of fusion. Fusion reactions were assembled as in Fig. 2. Where indicated, Sec17/18 (100 nM and 50 nM respectively) and ATP regenerating system (1 mM), were added to the RPLs 5 min. before the 5 min. baseline reading. Fusion was then initiated by adding 100 nM Sly1 at time= 0, then monitored for 30 min. Individual time points at 10 and 30 min. were taken for each reaction. Vertical bars show means ± S.E.M. for 3 independent experiments. P-values for selected 2-way ANOVA comparisons are indicated by horizontal bars (ns= P>.05, *P≤ 0.05, ** P≤0.01, ***P<0.001).

There was a delayed fusion signal with the R/Qb/Qc vs. Qa topology. Over the first 10 minutes virtually no fusion was detected; however, by 30 minutes there was a noticeable signal (Fig. 3B). It is important to note that heterotypic fusion generates post-fusion RPL populations that display all four SNAREs. Sec17/18:ATP may allow the products of slow or inefficient first-round fusion events to undergo additional rounds of more efficient fusion that do not reflect the activity of the starting SNARE topology. A delayed signal may suggest that inefficient fusion gradually creates a third population of RPLs bearing all four SNAREs, which can then fuse efficiently with at least one of the starting populations. We infer that this occurs with the R/Qb/Qc vs. Qa topology.

Sly1 binds the Qa SNARE Sed5 with such high affinity (KD < 1 nM) that under physiological conditions the complex probably does not dissociate, even during Sec18-mediated complex disassembly. (Demircioglu et al., 2014; Grabowski and Gallwitz, 1997; Khan et al., 2025). Indeed, a chimeric Sly1-Sed5 fusion protein can restore viability to a *sed5Δ* null mutant, demonstrating that Sed5 and Sly1 can function without dissociation (Gao and Banfield, 2020). We therefore asked whether pre-assembly of Sed5 and Sly1 prior to docking and Sec17/18 activity might alter the topological specificity of fusion. When this order of addition was tested for the R vs Qa/Qb/Qc topology, there was a small but reproducible increase in the initial rate of fusion (Duan et al., 2024a). To test whether the order of Sly1 addition alters topological requirements, we tested four additional topologies by adding Sly1 to Qa-bearing RPLs before priming by Sec17/18 or tethering (Fig. S4A). Early Sly1 addition did not stimulate fusion in the Qc vs R/ Qa/Qb topology (Fig. S4C). For both the R/Qc vs Qa/Qb and the R/Qb/Qc vs Qa topologies there was a decrease in the lag before fusion and a mild increase of total fusion stimulated (Fig. S4D-E). While there was a decrease in the lag before onset of fusion, there was not rapid fusion upon initiation of the reaction, as seen with the homotypic topology (Fig S4F). The lag period is still indicative. Early Sly1 addition does not appear to modify the topological constraints on fusion.

### Qb and Qc SNAREs transmembrane anchors are needed for efficient fusion

The yeast vacuole and presynaptic SNARE systems include one (Qc) or two (Qb and Qc) SNARE domains that lack a C-terminal transmembrane anchor. In chemically defined reconstitutions of yeast vacuole fusion (which included the HOPS tethering complex), fusion could be driven by soluble Qb and Qc SNAREs. We tested the limits of early secretory SNARE topology in reactions with Qb and Qc SNAREs that lack transmembrane anchors. The R and Qa SNAREs were always *in trans*, and when present, full length Qb or Qc SNAREs were *in cis* to the Qa. Soluble Qb or Qc SNARE cytoplasmic domains lacking transmembrane helices (Qb 1-222 and Qc 1-123), were added to reactions that contained the other full-length SNAREs, Sec17/18, ATP, and Sly1 (Fig. 4A). Even when soluble SNAREs were present at high concentrations (up to 4 µM), only modest fusion occurred, far below the level of fusion for the R vs Qa/Qb/Qc topology (Fig. 4B). Soluble Qb drove slightly more efficient fusion than the other configurations, and this mild stimulation was observed only in the presence of Sly1 and Sec17/18/ATP (Fig. 4C). These results provide further evidence that early secretory Qb and Qc SNAREs drive efficient, Sly1-dependent fusion when anchored in the same membrane as the Qa SNARE.

**Figure 4.**
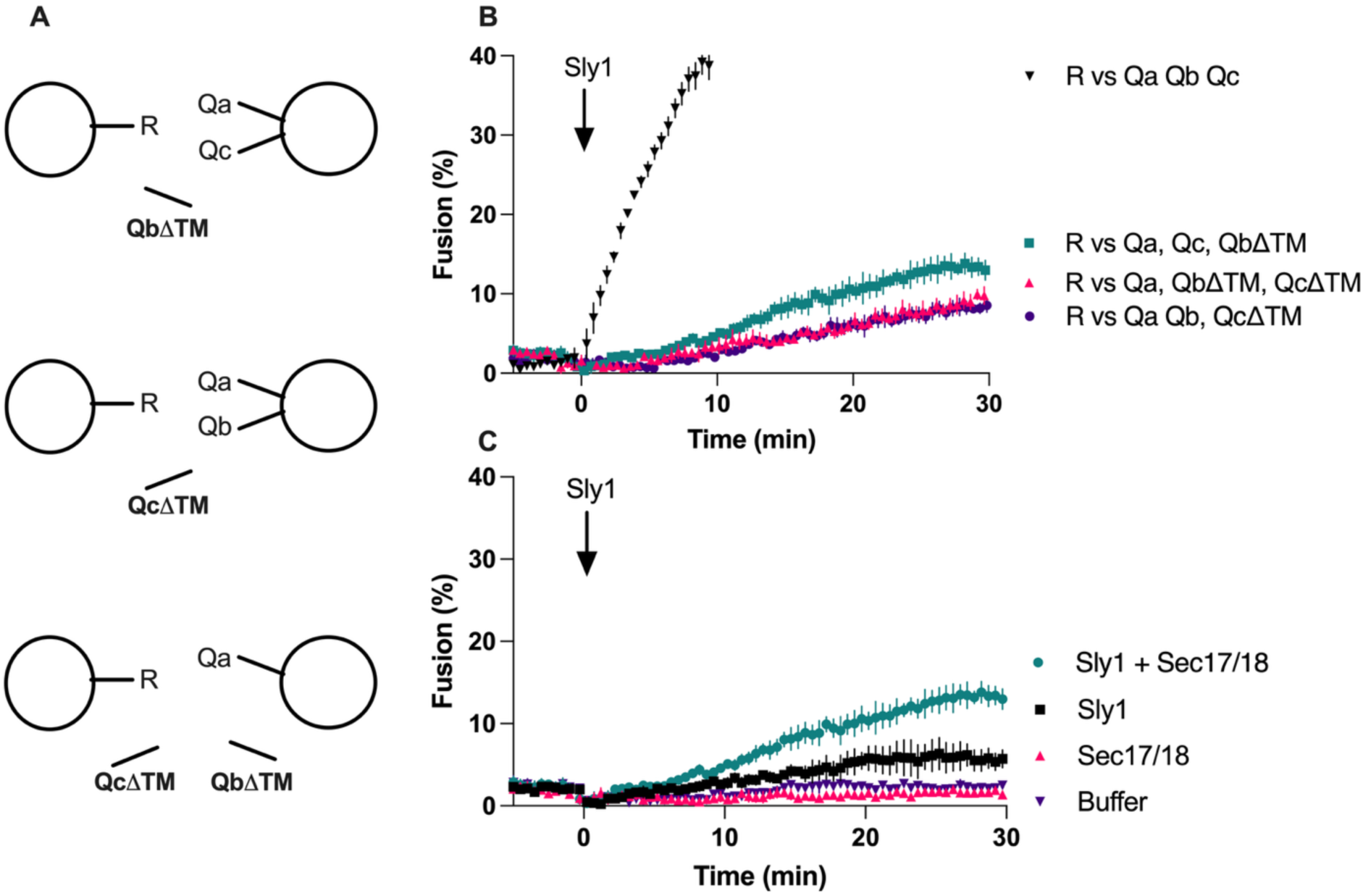
Qb and Qc SNAREs lacking membrane anchors do not drive efficient fusion. (**A**) Schematics show topologies used in soluble SNARE configurations. (**B, C**) Fusion reactions were set up and calibrated as described above with Sec17/18 (100 nM and 50 nM respectively) and ATP regenerating system (1 mM), and soluble SNAREs (4 µM each) were added prior to the 5 min baseline reading. Fusion was initiated by adding 100 nM Sly1 at time =0. (**C**) Fusion time course showing fusion conditions tested for R vs Qa in the presence of QbΔTM and QcΔTM (4 µM each). Points show means ± S.E.M. from 3 independent experiments.

### Sar1 and Sec23/24 are necessary and sufficient for COPII inhibition

With a better understanding of SNARE topological constraints in hand, we turned our attention to the machinery that forms the ER-Golgi carrier organelle. In cell extract experiments, when the full COPII coat (Sar1:GPPNP, Sec23/24, and Sec13/31) was locked onto the vesicle, fusion was blocked. This indicated that Sar1 GTP hydrolysis and at least partial COPII disassembly are prerequisites for fusion (Barlowe et al., 1994; Oka and Nakano, 1994). Because these early studies required the full COPII coat to form carriers, it was not possible to rigorously define the minimal set of COPII components that prevents fusion. Moreover, it was unclear whether additional factors might also be needed for COPII to block fusion. Although Sar1:GDP dissociation occurs rapidly, it is unclear whether the rest of the COPII complex dissociates at the same time, or if some subunits remain associated with the carrier (Antonny et al., 2001; Barlowe et al., 1994; Cai et al., 2007b; Lord et al., 2011; Sato and Nakano, 2005). It was therefore of interest to determine which subunits directly inhibit fusion.

To reactions containing RPLs bearing the R vs Qa/Qb/Qc SNAREs, we added purified COPII components to test how each component contributes to fusion inhibition (Fig. 5A-B). The Sec23/24 and Sec13/31 complexes were added to reactions at or slightly below their *in vivo* concentrations, while Sar1 was added to about twice its *in vivo* concentration (see Supplementary Table S1). This was done to increase Sar1:GTP levels, since the Sar1 nucleotide exchange factor Sec12 was not present in the reactions (Barlowe and Schekman, 1993).

**Figure 5.**
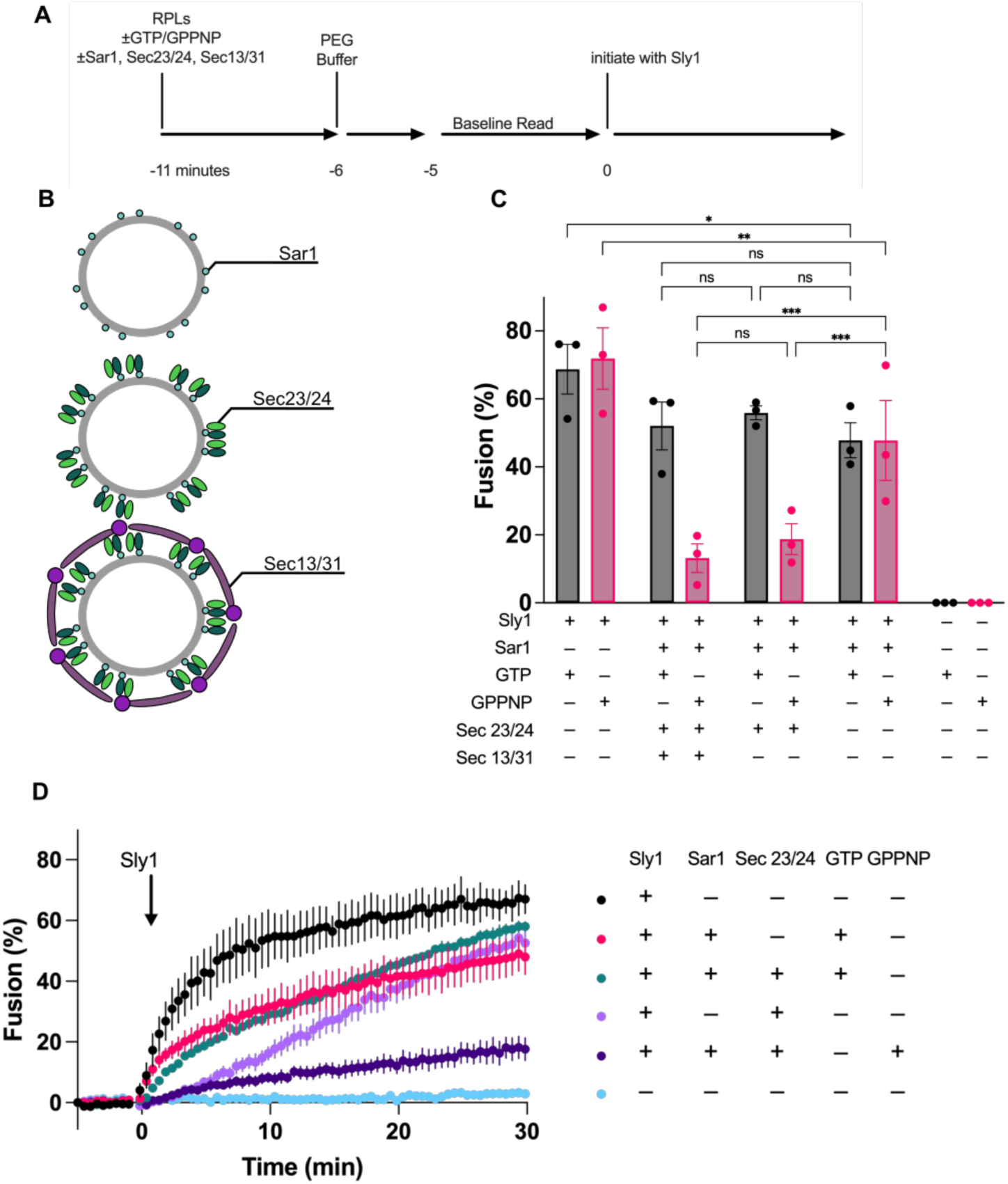
Sar1 and Sec23/24 are sufficient to inhibit fusion. Fusion assays were performed and calibrated as described above. Different combinations of COPII subunits were added to determine the minimal requirements for fusion. Sar1 was added to 2611 nM, Sec23/24 and mutant variants were added to 314 nM, and Sec13/31 was added to 407 nM. (**A**) Timeline showing the addition of each component to COPII fusion assays. (**B**) Schematic representation of the COPII layers and the tested conditions. (**C**) Fusion was measured after the addition of Sly1 at time=0 for 30 min. The last five minutes (t = 25 to 30 min) were averaged for each assay. Bars show means ±S.E.M. for 3 separate experiments. Grey bars show GTP conditions for each noted COPII protein combination; magenta bars show GPPNP conditions. P-values for selected 2-way ANOVA comparisons are indicated by horizontal bars (ns= P>.05, *P≤ 0.05, ** P≤0.01, ***P<0.001). (**D**) Fusion reactions with Sec23/24 and different nucleotide and GTPase conditions. The plot shows means ± S.E.M from 3 independent experiments.

Fusion was tested under both hydrolyzing (GTP) and non-hydrolyzing (GPPNP) conditions. As expected, in reactions with any combination of COPII protein and GTP we saw high levels of fusion both early (10 min) and near completion of the reaction (30 min). Sar1 GTP hydrolysis allows COPII to dissociate from the membrane for fusion to proceed. The initial rate of fusion, even in the presence of Sar1 alone, was lower than in reactions lacking COPII at 10 minutes (p=0.04) and 30 minutes (p=0.05). This decrease is observed across all three conditions. We suspect that membrane insertion of the Sar1 N-terminal helix partially opposes fusion (see Discussion).

In the presence of GPPNP, fusion was strongly impaired with all five COPII proteins present (Fig. 5C). These results recapitulate the original yeast extract studies and reveal that COPII by itself inhibits fusion, without any additional factors. We next tested combinations of the COPII components for inhibition of fusion. In reactions containing Sar1:GPPNP and Sec23/24, but lacking Sec13/31, fusion was inhibited as completely as in the presence of all five COPII components. Thus, the inner coat complex (Sar1:GPPNP and Sec23/24) is sufficient to block fusion. The Sec13/31 outer shell does not contribute to inhibition (Fig. 5C).

In the yeast extract system, higher concentrations of COPII, Sec23/24 complex, or recombinant GST fusions to Sec23 or Sec24 inhibit COPII carrier tethering and fusion, even under GTP-hydrolyzing conditions (Barlowe, 1997; Cai et al., 2007b). We therefore tested whether Sec23/24 could inhibit fusion when Sar1 was not present. There was substantial inhibition early in the reaction, but this inhibition diminished over time (Fig. 5D). Sec23/24 binds membranes through its affinities for Sar1:GTP, the membrane itself, and cargo proteins including SNAREs. The results suggest that, even without Sar1, Sec23/24 heterodimer can partially inhibit fusion. This is consistent with experiments where Sar1 is released rapidly during carrier formation but the remaining COPII subunits dissociate more slowly (Barlowe, 1997; Cai et al., 2007b). Perhaps unexpectedly, Sec23/24 alone appears to inhibit fusion almost as completely as Sar1:GPPNP and Sec23/24 for the initial 10 minutes of the reaction, but by 30 minutes behaves more like Sar1:GTP and Sec23/24 (Fig. 5D). We conclude that Sec23/24 is primarily responsible for fusion inhibition, while Sar1:GPPNP causes stable Sec23/24 association and prolonged inhibition of fusion. The early inhibition observed here may indicate that, *in vivo*, active mechanisms promote fusion by stripping Sec23/24 from the carrier. Candidate factors for such an activity include the tethering factor Grh1, casein kinase I, and, in multicellular organisms, TFG (Behnia et al., 2007; Hanna et al., 2017; Humphreys et al., 2021; Lord et al., 2011).

### Sec23/24 binding interactions with SNAREs are necessary for inhibition

Does COPII act as a relatively nonspecific “steric mesh” that prevents the membranes from coming into close contact, or does inhibition depend mainly on specific regulatory coat-SNARE interactions? Stereospecific interactions between Sec23/24 and SNAREs package SNAREs at ER exit sites, ensuring that COPII carriers are competent to fuse with target membranes. However, COPII-SNARE interactions might directly regulate the ability of SNAREs to mediate fusion. There are three distinct cargo binding sites on Sec24, each recruiting different SNAREs.

Missense mutations in the Sec24 cargo binding sites disrupt interactions with SNAREs and other cargo. Two specific point mutants, L616W and W897A, are of special interest. Like wild type Sec24, these mutants assemble into stable lattices and do not impair COPII carrier formation. These mutants were originally identified through defects in their ability to package SNAREs into vesicles. In our experiments, the SNAREs are already present on the RPLs, so the cargo binding mutants can be used to test whether Sec24-SNARE binding interactions are important for COPII inhibition at the fusion stage. Sec24 L616W disrupts the B-site, impairing its interaction with the Qc and R SNAREs, and *in vivo*, cannot support cell viability (Miller et al., 2003). Sec24 W897A disrupts the A-site, which mediates Qa SNARE binding. *In vivo*, the W897A mutation is lethal if the partially redundant Sec24 paralog Sfb2 is deleted (Miller et al., 2005). In fusion assays with GPPNP, the B-site mutant Sec23/Sec24 (Sec24 L616W) inhibited fusion less strongly than the wild type, and similarly to Sar1-only conditions (Fig. 6C-D). In marked contrast, mutation of the Qa-only A-site pocket (Sec24 W897A) did not relieve inhibition in the presence of GPPNP. Moreover, in reactions with GTP and the A-site mutant, fusion was somewhat slower than with wild type Sec23/24. (Fig. 6A-D). As we had found above that wildtype Sec23/24 was capable of transiently inhibiting fusion without Sar1, we tested whether this inhibition required interactions with SNAREs. We found that Sec23/24 containing either SNARE binding mutation had little or no inhibitory effect. (Fig. 6E-F). We conclude that SNARE sequestration within the Sec24 B-site is the primary mechanism through which COPII inhibits fusion.

**Figure 6.**
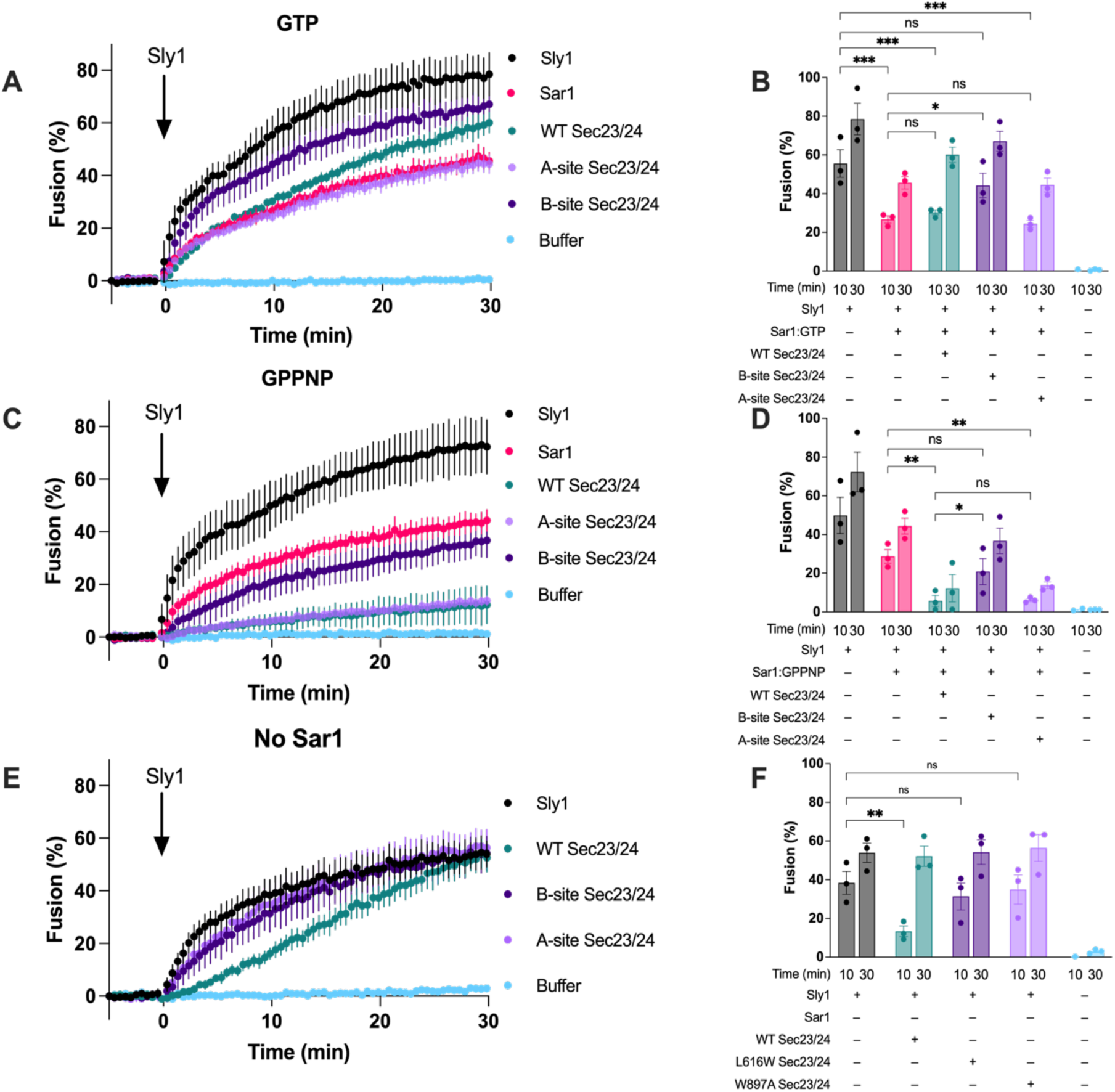
Sec24-SNARE interactions regulate fusion. Fusion assays were performed as previously described. Reactions included permutations of nucleotides, Sar1, and Sec23/24 wild-type, or Sec23 A-site (W897A) or B-site (L616W) mutant protein, added to the same concentrations as in Fig. 5. Fusion was measured after addition of Sly1 at time = 0. Sar1 was added to 2611 nM, Sec23/24 and mutant variants were added to 314 nM, and Sec13/31 was added to 407 nM. (**A** & **B**) Reactions contained GTP. (**C** & **D**) Reactions contained GPPNP. (**E** & **F**) Reactions contained no nucleotide and no Sar1. (**A**, **C**, & **E**) Fusion assays initiated with 100 nM Sly1 at time=0. Each point represents the means of 3 independent reactions ± S.E.M. (**B, D,** & **F**) Measurements at 10 and 30 min (±0.5 min) were taken from the time courses shown in panels A, B, or E respectively. Bars represent means with error bars representing ± S.E.M. P-values for selected 2way ANOVA comparisons are indicated by horizontal bars (ns= P>.05, *P≤ 0.05, ** P≤0.01, ***P<0.001).

## Discussion

Our results argue that ER-Golgi fusion requires the R SNARE oppose the Qa and Qb SNAREs. The Qc SNARE may act on either membrane, but is most efficient *in cis* to the Qa and Qb SNAREs. As discussed below, strong evidence indicates that the Qa SNARE Sed5 functions on the Golgi acceptor membrane. Thus, the R SNARE Sec22 is the primary fusogenic SNARE on the COPII carrier. Consistent with these findings, we find that Sec24 binding to R and Qc SNAREs is the primary mechanism through which COPII inhibits fusion. This paints a satisfying picture: COPII first ensures that the R (and perhaps Qc) SNAREs required for fusion are packaged into the carrier. It then holds them ready but inactive until the coat’s inner shell has been shed.

Little or no basal fusion occurs in the absence of Sly1 with any SNARE topology, consistent with the lethality of *sly1*Δ mutants. In the presence of Sec17/18 and ATP, fusion is strongly stimulated when all four SNAREs are present on both membranes, since disassembly of *cis*-SNARE bundles is required for fusion in the homotypic configuration. Even when all SNAREs are present on both membranes, it is reasonable to assume that productive fusion reactions employ the R SNARE in *trans* to Qa/Qb/Qc. Thus, the intrinsic properties of SNAREs and the SM act together to enforce topological constraints on fusion. The requirement for R and Qa SNAREs to operate *in trans* is consistent with crystallographic, biophysical, and modeling studies. In contrast, it is not yet possible to explain the requirement for Qb and Qc SNAREs to operate *in cis* to the Qa.

Previously, Rothman’s group examined basal ER-Golgi fusion in the absence of the SM Sly1, and identified a unique functional topology: Qc opposite Qa, Qb, and R (McNew et al., 2000a; Parlati et al., 2000; Parlati et al., 2002). This discrepancy between our experiments results from the presence or absence of Sly1, the fusion readouts used, and perhaps other differences in methodology. We primarily used content mixing signals while the Rothman experiments relied solely on lipid mixing. Importantly, we observed very slow but reproducible lipid mixing with the Qc vs. Qa/Qb/R topology, but there was no stimulation by Sly1 with this topology (Fig. S3). Additionally, our studies used different lipid-to-protein ratios, different lipid compositions, and different lipid mixing reporters. *In vivo*, all SNARE-mediated fusion requires an SM (Sudhof and Rothman, 2009).

Experiments with cell extracts demonstrated that Sed5 (Qa) and Sly1 must act at the target membrane, and are not needed on the COPII carrier (Cao and Barlowe, 2000; Miller et al., 2005). Cao and Barlowe further suggested that Bos1 (Qb) and Bet1 (Qc) function only on the COPII carrier, along with Sec22 (R) (Liu and Barlowe, 2002). In contrast, our experiments suggest that fusion is most efficient when all three Q SNAREs are *in cis*. Fusion with the R/Qb/Qc vs Qa topology was slow and exhibited a long lag after initiation. A stronger fusion signal was observed when Sly1 was preincubated with the Qa SNARE RPLs, but preincubation did not overcome the long lag (Fig. S4E). We conclude that the R/Qb/Qc vs Qa topology does not efficiently catalyze fusion. The basis for the discrepancy between our results and those of Cao and Barlowe is unclear. One possibility is that additional factors present in the cell extract system could influence SNARE topology requirements. For example, the use of temperature-sensitive SNARE mutants may perturb interactions with COPII or tethering factors upstream of *trans* SNARE complex assembly. Interactions between the Uso1/p115 tether and R, Qb and Qc SNAREs were recently established (Bravo-Plaza, et. al., 2023). Thus, it is possible that these SNAREs have functions in tethering distinct from their roles in direct catalysis of fusion.

Previous chemically defined reconstitutions of fusion yielded divergent results with respect to the specificity and selectivity of SNARE topological restriction. These studies were mostly done in the absence of SMs (Fukuda et al., 2000; Furukawa and Mima, 2014; McNew et al., 2000a; Parlati et al., 2000; Parlati et al., 2002; Zwilling et al., 2007). In the yeast vacuole system, the HOPS complex contains the SM Vps33, but HOPS contains five other subunits, some of which bind SNAREs (Dulubova et al., 2001; Lobingier and Merz, 2012; Song and Wickner, 2017; Wickner et al., 2023). In the vacuole system the Qc SNARE is soluble. Surprisingly, no published experiments in this highly characterized system have directly compared fusion with native (non-mutant) SNAREs in the three possible arrangements of R, Qa, and Qb. To our knowledge, in fact, only one previous study, from Shen’s laboratory, has systematically explored SNARE topology in the presence of an SM and without the potentially confounding presence of other tethering factors (Rathore et al., 2011). In their experiments, chimeric neuronal SNARES bearing Qb and Qc SNARE domains were fused to C-terminal transmembrane domains. Fusion was totally dependent on Munc18-1, and efficient only with the R vs. Qa/Qb/Qc topology. Although the Shen group’s system is synthetic, their results suggest that underlying mechanisms that enforce topology in the early secretory pathway may be conserved across SM-SNARE systems.

Yeast cells lacking the R-SNARE Sec22 are thermosensitive and grow slowly, but viable (Dascher et al., 1991; Liu and Barlowe, 2002; Ossig et al., 1991). In these cells the R-SNARE Ykt6 substitutes for Sec22 (Liu and Barlowe, 2002). Ykt6 is promiscuous, operating in other Golgi fusion events as well as vesicle and autophagosome fusion with lysosomes (Bas et al., 2018; Gao et al., 2018; Kweon et al., 2003; Matsui et al., 2018; McNew et al., 1997; Takáts et al., 2018; Tateishi et al., 2025; Watanabe et al., 2024). Ykt6 lacks a transmembrane anchor, instead associating with membranes through farnesyl and geranylgeranyl moieties appended Cys-Cys motif at its C-terminus (McNew et al., 1997; Sakata et al., 2021; Shirakawa et al., 2020; Tateishi et al., 2025). If Ykt6, like Sec22, acts *in trans* to Qa, Qb, and Qc SNARES, then it must drive fusion with no α-helical transmembrane anchor on one membrane. However, when either the Qa or R transmembrane segment of a neuronal SNARE complex was replaced with prenyl anchors in a minimal reaction system, anchors longer than the geranylgeranyl group were required to initiate lipid mixing (McNew et al., 2000b). It will be illuminating to examine the mechanisms of Ykt6-mediated fusion in chemically defined systems that incorporate SMs and other cofactors.

COPII is a paradigm for understanding how coats couple carrier formation to downstream docking and fusion events. In crude extract systems the complete COPII complex prevents fusion (Barlowe et al., 1994; Oka and Nakano, 1994). Our experiments identify Sar1 and Sec23/24 as both necessary and sufficient for this inhibition (Fig. 5C). Sec23/24 partially inhibited fusion without Sar1. Binding between the SNAREs and Sec24 is critical for SNARE packaging into vesicles with disruption of the SNARE-COPII interaction leading to absence of SNAREs in COPII vesicles. Distinct sites bind different SNAREs (Miller et al., 2003, Miller et al. 2005, Mossessova et al., 2003). The Sec24 B mutant employed here, L616W, most strongly impairs packaging of Sec22 (R) and Bet1 (Qc). We found that L616W allows fusion co occur in the presence of GPPNP-locked Sar1. Thus, the Sec24 B site, in addition to cargo packaging, specifically prevents fusion by sequestering SNAREs. Consistent with this model, in cell-free extracts recombinant Sec23 inhibited tethering and subsequent fusion, while excess recombinant Sec24 inhibited fusion without disrupting tethering (Barlowe, 1997; Cai et al., 2007b). The assembled COPII mesh may also prevent fusion by preventing membrane approach, but it is not entirely clear how complete, patchy, or dynamic the COPII coat is *in vivo*, particularly in the presence of GTP rather than its non-hydrolyzable analogs.

Unexpectedly, Sar1 partially inhibited fusion in the absence of other COPII subunits. Penetration of the membrane cytoplasmic leaflet by Sar1’s amphipathic N-terminal helix generates positive membrane curvature. This curvature is hypothesized to promote COPII carrier formation and carrier scission from the ER exit site (Kozlov and Taraska, 2023; Lee et al., 2005; Paul et al., 2023; Van der Verren and Zanetti, 2023). In contrast, lysolipids added to the cytoplasmic leaflet promote positive curvature and fission, and potently inhibit fusion (Chernomordik et al., 1993; Günther-Ausborn et al., 1995; Melia et al., 2006; Reese and Mayer, 2005; Schwartz and Merz, 2009; Vogel et al., 1993). COPII carriers budded from native ER membranes are enriched in lysolipids (Melero et al., 2018). As COPII carriers form in the presence of Sar1, GTP hydrolysis and Sar1 dissociation may relieve compression within the outer leaflet, thereby relaxing positive membrane curvature and allowing fusogenic membrane structures to form. In an alternative model, Sar1:GTP directly engages SNARE proteins and places them in an inactive state. The interplay between Sar1 and SNARE-mediated fusion will require additional scrutiny.

The directionality of vesicle transport is regulated by sequential interactions of SNAREs with coat proteins, other SNAREs, and chaperones. We have shown that a specific interaction between Sec24 and SNAREs prevents fusion, ensuring that the vesicle cannot back fuse with the ER, and maintaining functional separation from other SNARE-containing membranes. Topological restrictions further confer fusion specificity to membranes based on the presence or absence of specific SNAREs and SM proteins. The requirements for fusion we have identified support a model of strict sequential and topological regulation throughout the transport process, starting with SNARE packaging and ending with the fusion event itself.

## Materials and Methods

### Protein purification

Full-length SNAREs and Sly1 were purified as described (Duan et al., 2024b). Soluble SNAREs were purified in the same manner as Sly1. Sar1 was purified as described (Shimoni and Schekman, 2002). Sec23/24 complexes and Sec13/31 complexes were purified as described (Stancheva et al., 2020). Plasmids used for protein expression and purification are listed in Table S2.

### Preparation of RPLs

Reconstituted proteoliposomes were made as previously described (Duan et al., 2024b) with minor modifications. Lipid compositions were adjusted such that each population was identical, with the exception of the fluorescent reporter lipid. Table S3 describes the liposome compositions and vendor sources.

### Fusion assays

Fusion was assayed and calibrated as described (Duan et al., 2024b). For reactions including RPLs, unlabeled streptavidin and soluble SNAREs were pre-mixed before the addition to a final concentration of 3% PEG, Sec17/18 (100 nM, 50 nM, or omitted), ATP regenerating mixture to 1 mM, and additional buffer. For reactions containing COPII proteins, Sar1 was added at 2611 nM, Sec23/24 and mutant variants were added at 314 nM, Sec13/31 was added at 407 nM. These reactions were immediately transferred to the plate reader and baseline fluorescence readings were acquired. Fusion was initiated by adding Sly1 (100 nM). Content mixing calibration was carried out for each RPL pair tested. 100% was determined by the total possible content mixing signal when both liposome populations were lysed with detergent in the absence of unlabeled competing streptavidin. Zero percent fusion was calibrated by taking baseline fluorescence measurements of reactions containing RPLs but not soluble cofactors.

### AlphaFold3 modeling

Models of SNARE and SM interactions were modeled using the Google AlphaFold3 server. SNAREs were truncated from the N-terminus to their SNARE domain. R and Qa SNAREs maintained the remainder of the protein which Qb and Qc SNAREs were truncated at their –3 layer to show a partially fused complex. Modeling parameters were set to defaults.

### Estimation of *in vivo* protein concentrations

*In vivo* protein copy numbers and concentrations (Supplementary Table S1) were estimated as before (Duan et al., 2024b). From https://www.yeastgenome.org, curated Protein Abundance data for each protein were filtered to include only estimates for cells grown in YEPD media, and with no other specified treatments (drugs, *etc.*). Geometric means of the filtered copy number data are reported, to decrease sensitivity to high-count outlier measurements. Our estimates are in good agreement with comparable estimates by others (Ho et al., 2018). The volume of a haploid yeast cell is assumed to be ∼67 fL, based on light microscopy and X-ray tomography measurements (Chan et al., 2016; Chan and Marshall, 2014; Uchida et al., 2011). The volume of accessible cytosol (cytoplasm minus major organelles including nucleus, ER, vacuole, mitochondria etc.) is taken as 75% of this volume, or 50 fL, similar to the fraction estimated by Uchida et al.

## ACKNOWLEDGMENTS

We thank M. Munson, J. McNew, R. Baker, and S. Hoppins for insightful discussions over the course of this work, and C. Barlowe for sharing reagents and strains. We thank the referees for constructive suggestions that improved the study.

## Supplementary Figures

**Supplementary Figure 1.**
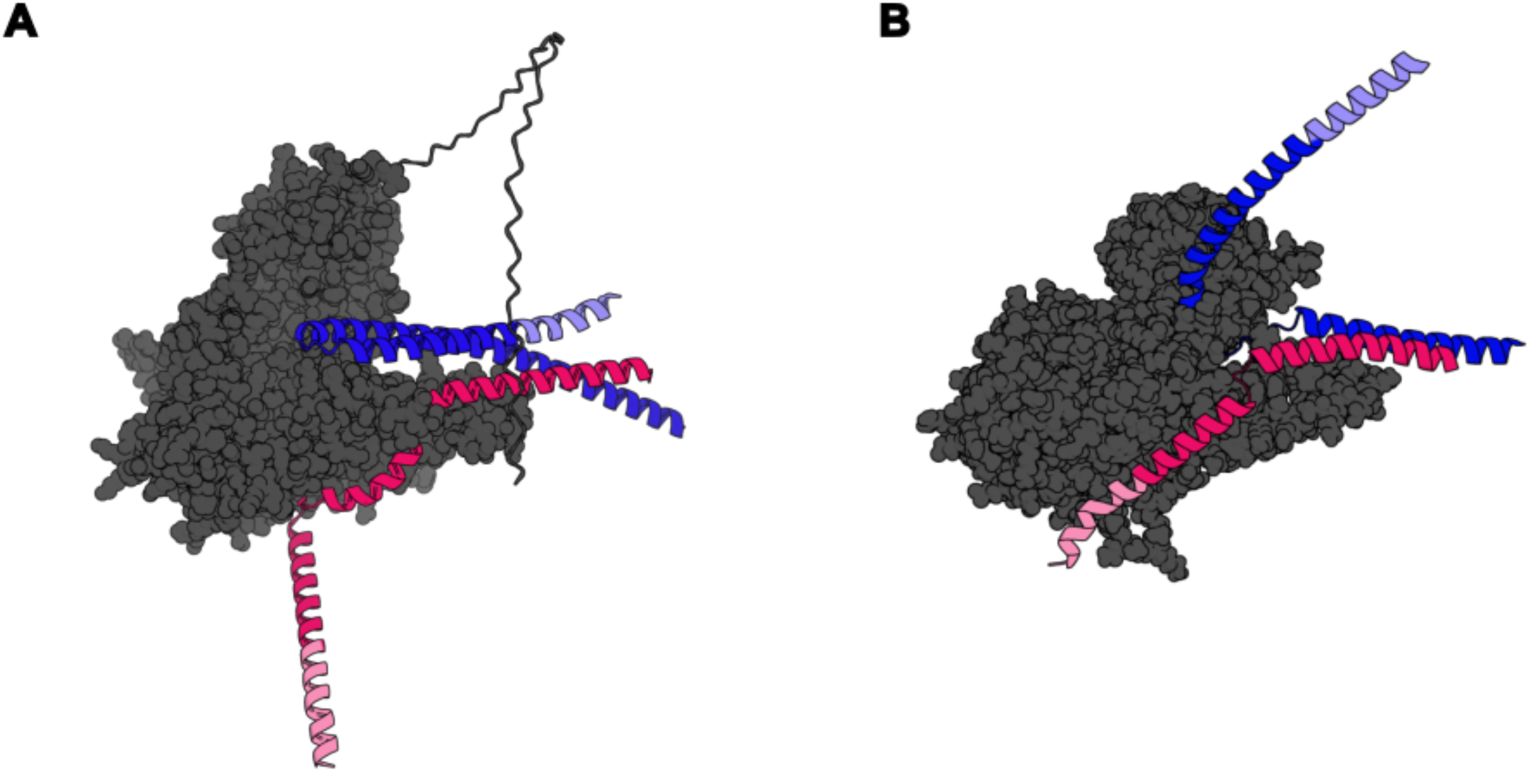
AlphaFold3 modeling of SMs and cognate Qa and R SNAREs. (**A**) Sec1 (SM) with Sso1 (Qa) and Snc2 (R). Sec1 is shown in grey with residues 670-724 shown as ribbon for clarity. Sso1 (residues 183-289), is colored blue with the TMD in light blue (274-289). Snc2 (residues 20-115), is colored red with the TMD in pink (98-115). (**B**) Vps45 (SM) with Tlg2 (Qa) and Ykt6 (R) AlphaFold 2 model. Vps45 is shown in grey. Tlg2 (residues 243-336) is colored blue, with its TMD shown in light blue (318-336). Ykt6 (residues 138-199) is colored red with its membrane-proximal region (186-199) in pink. The C-terminal prenyl lipids that anchor Ykt6 to the membrane are not modeled.

**Supplementary Figure 2.**
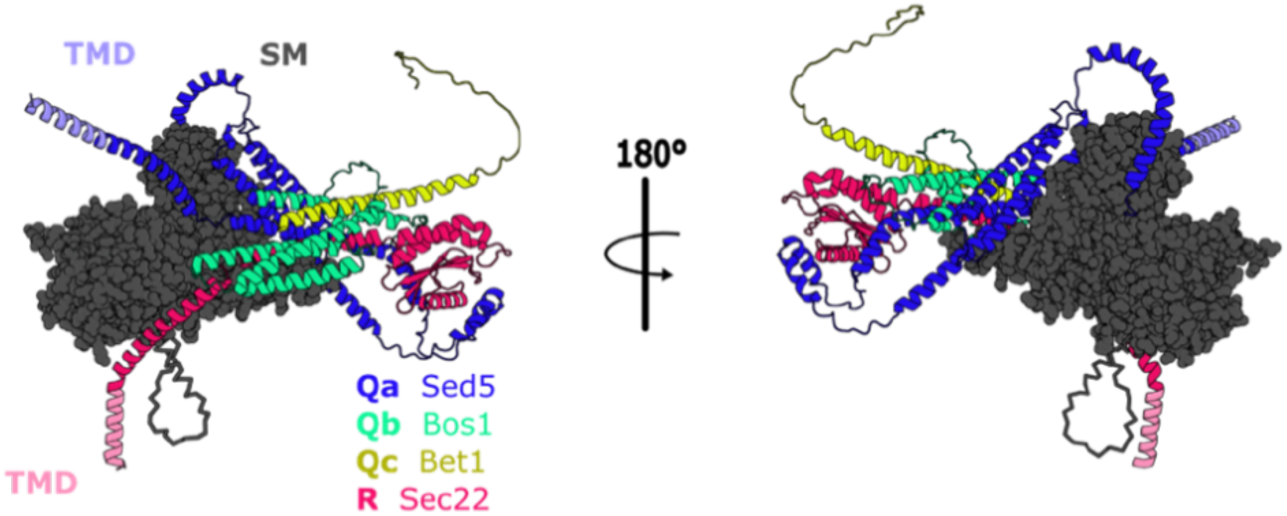
The N-terminal regulatory domains of SNAREs are not predicted to modify their SNARE templating interactions with Sly1. AlphaFold3 models of Sly1 with full length R (Sec22) and Qa (Sed5) as well as the Qb (Bos1) (1-182), and Qc (Bet1) (1-78) to their –3 layer. Modeling includes the N-terminal domains depicting additional interactions between the SNAREs. The absence or presence of the Sec22 N-terminal longin domain does not dramatically alter the positions of the Qa or R SNARE domains on Sly1 domain 3a, or the binding of Sec22 Phe186 in conserved pocket on Sly1 (also seen in published and deposited but unpublished Vps33-Nyv1 crystal structures: PDB 5BV0, 9ZCS, 9BZE, 9ZCT).

**Supplementary Figure 3.**
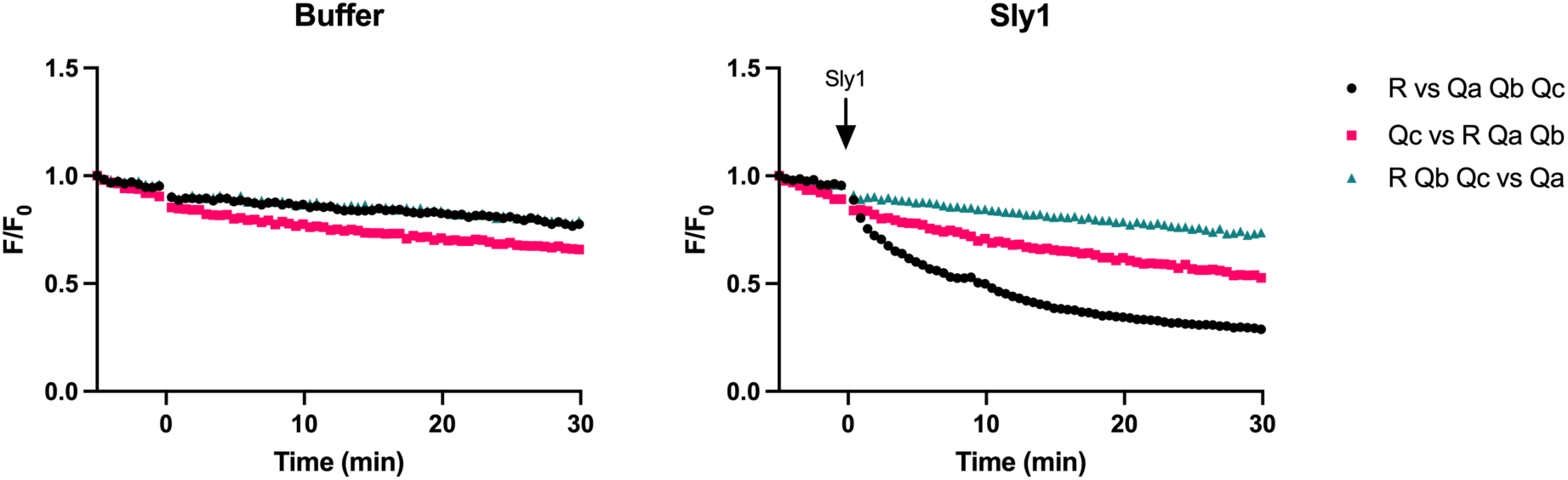
Lipid mixing with representitive topologies. Representative lipid mixing signals in the absence (**A**) and presence (**B**) of Sly1. Raw fluorescence was normalized to the starting signal for each population. Note that in panel A, the traces for R vs. Qa Qb Qc, and R Qb Qc vs. Qa overlap so closely that they are not distinguishable.

**Supplementary Figure 4.**
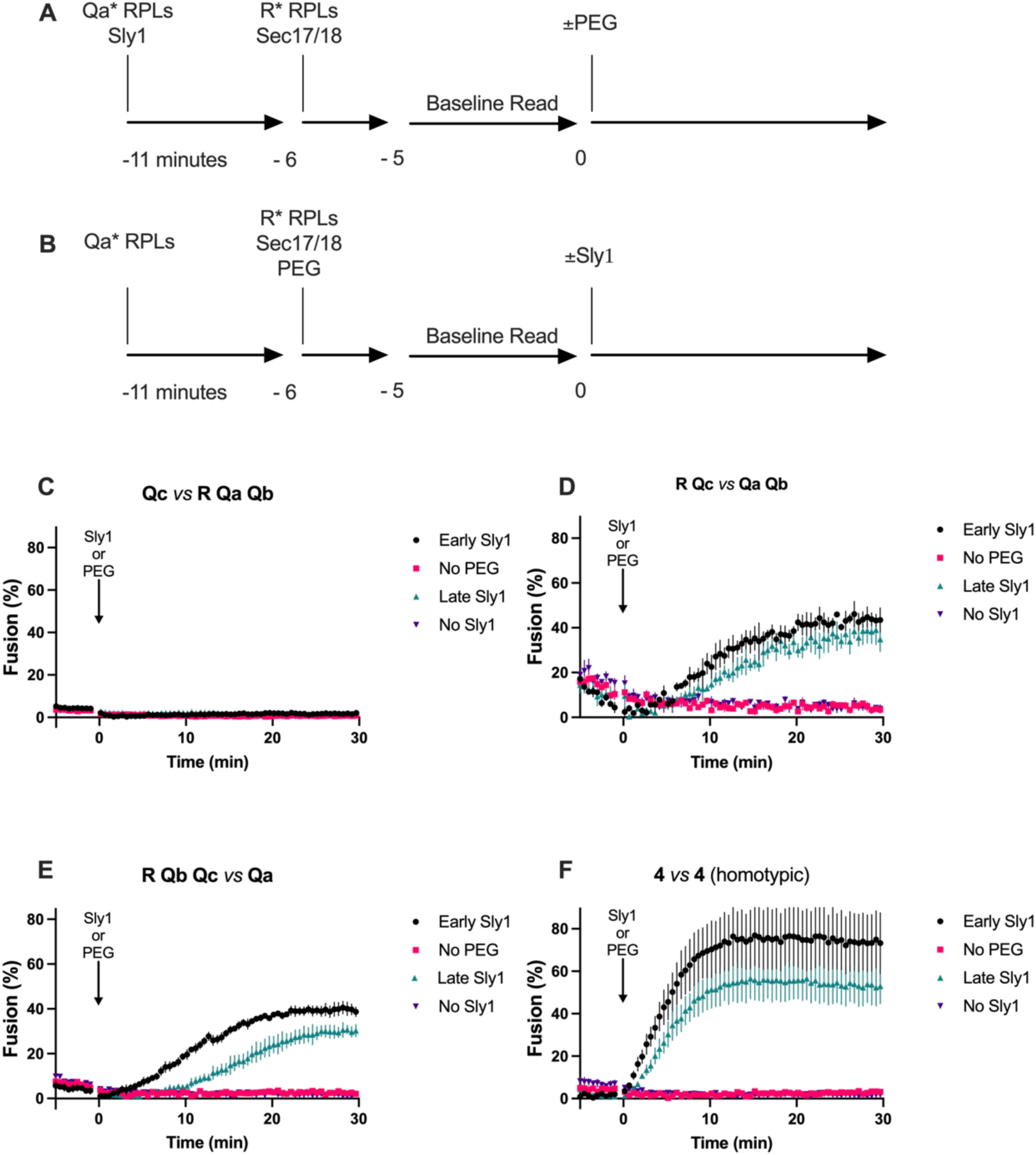
Effects of Sly1 order-of-addition. Order of addition experiments to determine if Sly1 and Qa interactions can additionally stimulate fusion before the addition of the other RPL population. (**A** and **B)** Schematic representations of the order of addition of each component to the fusion assays in this figure. The top panel (A) shows early Sly1 addition, while the bottom panel shows late sly1 addition. Qa* represents the RPL population bearing the Qa SNARE (with or without other SNAREs, depending on the topology tested). For the homotypic RPL set only one population was added at this time point. Sec17/18 and ATP were added in all experiments. Experiments with Sly1 added early to the Qa* population (A) were initiated with PEG at time=0. This order of addition was used for the black (circle) and pink (square) traces in the below graphs. Experiments initiated with Sly1 added late (B) are similar to the order-of-addition used all other topology experiments in this study. This order of addition was used for the teal (triangle) and purple (inverted triangle) traces. (**C-F**) Time course traces, each for a different SNARE topology, showing the kinetic effects of adding Sly1 to the Qa* population.

## Supplementary Tables

**Supplementary Table 1.** Comparison of estimated *in vivo* protein concentrations versus concentrations used in this study’s fusion reconstitution experiments.

|  |  |  |  |  |  |
| --- | --- | --- | --- | --- | --- |
| | Cytosol volume, $\mu\text{m}^3$ (fL) | Copy number, molecules/cell | Concentration in or exposed to cytosol, nM | Concentration in RPL fusion assays in this study, nM | Notes |
| **Sec17** | 50 | 11,364 | **378** | **100** | SNARE adapter for Sec18 |
| **Sec18** | 50 | 8,115 | **270** | **50** | SNARE disassembly ATPase |
| **Sly1** | 50 | 6,380 | **212** | **100** | SM family SNARE chaperone |
| **Sed5** | 50 | 2,728 | **91** | **$\leq 200$** | Golgi Qa-SNARE; Sly1 client |
| **Bos1** | 50 | 2,870 | **95** | **$\leq 200$** | Golgi Qb-SNARE |
| **Bet1** | 50 | 3,772 | **125** | **$\leq 200$** | Golgi Qc-SNARE |
| **Sec22** | 50 | 3,288 | **109** | **$\leq 200$** | Golgi/ER R-SNARE |
| **Sar1** | 50 | 22,568 | **750** | **1600** | COPII small GTPase |
| **Sec23** | 50 | 14,568 | **484** | **300** | COPII inner shell |
| **Sec24** | 50 | 8,873 | **295** | **300** | COPII inner shell |
| **Sfb2** | 50 | 3,615 | **120** | - | COPII inner shell (Sec24 paralog) |
| **Sfb3** | 50 | 7,835 | **260** | - | COPII inner shell (Sec24 paralog) |
| **Sec13** | 50 | 19,617 | **652** | **400** | COPII outer shell & nuclear pore |
| **Sec31** | 50 | 14,493 | **481** | **400** | COPII outer shell |

**Supplementary Table 2.** Plasmids used in this study.

| Supplementary Table 2. Plasmids used in this study |  |  |
| --- | --- | --- |
| plasmid ID | Description | Source/reference |
| *E. coli* SNARE expression plasmids |  |  |
| AMP1792 | pET-30::His<sub>6</sub>-(3C)-Sed5 | (Furukawa and Mima, 2014) |
| AMP2020 | pET-30:: His<sub>6</sub>-(3C)-Bos1(Q153D) | (Furukawa and Mima, 2014), but modified per (Duan et al., 2024) |
| AMP1794 | pET-30:: His<sub>6</sub>-(3C)-Bet1 (with corrected missense mutation) | (Furukawa and Mima, 2014) but corrected as (Duan et al., 2024) |
| AMP1795 | pET-41::GST- His<sub>6</sub>-(3C)-Sec22 | (Furukawa and Mima, 2014) |
| AMP1977 | pET-30:: His<sub>6</sub>-(3C)-Bos1(Q153D)(1-222) | This Study |
| AMP1976 | pET-30:: His<sub>6</sub>-(3C)-Bet1(1-123) | This Study |
| *E. coli* SNARE chaperone expression plasmids |  |  |
| AMP1547 | pTYB12::intein-CBD-Sec17 | (Schwartz and Merz, 2009) |
| AMP77 | pQE9:: His<sub>6</sub>-SEC18 | (Haas and Wickner 1996) |
| *E. coli* Sly1 expression plasmids |  |  |
| AMP1649 (pBL51) | pHIS-Parallel1:: His<sub>6</sub>-(TEV)-Sly1 | (Lobingier and Merz, 2014) |
| AMP1651 | pHIS-Parallel1:: His<sub>6</sub>-(TEV)-Sly1-20 | (Duan et al., 2024) |
| *E. coli* miscellaneous expression plasmids |  |  |
| AMP1881 | pET-49::GST- His<sub>6</sub>-(3C) | (Duan et al., 2024) |
| AMP2019 | pET-30:: His<sub>8</sub>-HRV3C(Protease) | (Duan et al., 2024) |
| AMP2016 | pET-49::GST- His<sub>6</sub>-(Thrombin)-HRV3C(Protease) | (Duan et al., 2024) |

**Supplementary Table 3.** SNARE RPL lipid compositions used in this study.

| Supplementary Table 3. SNARE RPL lipid compositions used in this study |  |  |  |  |  |  |
| --- | --- | --- | --- | --- | --- | --- |
| ER mimetic RPLs |  |  |  | Golgi mimetic RPLs |  |  |
| Lipid | Vendor | mol% |  | Lipid | Vendor | mol% |
| 18:1 (Δ9-Cis) PC | Avanti 850375 | 35.0% |  | 18:1 (Δ9-Cis) PC | Avanti 850375 | 35.0% |
| 16:0-18:1 PE | Avanti 850757 | 16.0% |  | 16:0-18:1 PE | Avanti 850757 | 14.0% |
| Soy PI | Avanti 840044 | 6.3% |  | Soy PI | Avanti 840044 | 6.3% |
| 16:0-18:1 PS | Avanti 840034 | 5.6% |  | 16:0-18:1 PS | Avanti 840034 | 5.6% |
| 16:0-18:1 PA | Avanti 840857 | 3.5% |  | 16:0-18:1 PA | Avanti 840857 | 3.5% |
| 18:1 DG | Avanti 800811 | 1.4% |  | 18:1 DG | Avanti 800811 | 1.4% |
| Brain PI(4)P | Avanti 840045 | 2.0% |  | Brain PI(4)P | Avanti 840045 | 2.0% |
| Ergosterol | Sigma 45480 | 30.0% |  | Ergosterol | Sigma 45480 | 30.0% |
| 16:0 Marina Blue™ DHPE | Invitrogen M12652 | 0.2% |  | 16:0 NBD-PE | Invitrogen N360 | 2.2% |

## Notes

### Competing Interest Statement

The authors have declared no competing interest.

### Summary of Updates

This version includes clarifications to the text, additional statistical analyses, new figures and figure panels, and an additional table.

### Funding

- National Institutes of Health: R01 GM077349, R01 GM130644
- Wellcome Trust: 225216/Z/22/Z
- UK Research and Innovation: MC_UP_1201/10

